# Network Curvature as a Hallmark of Brain Structural Connectivity

**DOI:** 10.1101/162875

**Authors:** Hamza Farooq, Yongxin Chen, Tryphon T. Georgiou, Allen Tannenbaum, Christophe Lenglet

## Abstract

Studies show that while brain networks are remarkably robust to a variety of adverse events, such as injuries and lesions due to accidents or disease, they may be fragile when the disturbance takes place in specific locations. This seems to be the case for diseases in which accumulated changes in network topology dramatically affect certain sensitive areas. To this end, previous attempts have been made to quantify robustness and fragility of brain functionality in two broadly defined ways: (i) utilizing model-based techniques to predict lesion effects, and (ii) studying empirical effects from brain lesions due to injury or disease. Both directions aim at assessing functional connectivity changes resulting from structural network variations. In the present work, we follow a more geometric viewpoint that is based on a notion of curvature of networks, the so-called Ollivier-Ricci curvature. A similar approach has been used in recent studies to quantify financial market robustness as well as to differentiate biological networks corresponding to cancer cells from normal cells. The same notion of curvature, defined at the node level for brain networks obtained from MRI data, may help identify and characterize the effects of diseases on specific brain regions. In the present paper, we apply the Ollivier-Ricci curvature to brain structural networks to: i) Demonstrate its unique ability to identify robust (or fragile) brain regions in healthy subjects. We compare our results to previously published work which identified a unique set of regions (called *structural core*) of the human cerebral cortex. This novel characterization of brain networks, complementary to measures such as degree, strength, clustering or efficiency, may be particularly useful to detect and monitor candidate areas for targeting by surgery (e.g. deep brain stimulation) or pharmaco-therapeutic agents; ii) Illustrate the power our curvature-derived measures to track changes in brain connectivity with healthy development/aging and; iii) Detect changes in brain structural connectivity in people with Autism Spectrum Disorders (ASD) which are in agreement with previous morphometric MRI studies.

## 1. Introduction

This paper describes a geometric network-theoretic approach to study brain structural connectivity. As is well-known, comprehensive structural connectivity of the human brain, the so-called ***connectome*** [1, 2], is not available to date despite recent progress using neuroimaging such as the Human Connectome Project (HCP) [3–6]. Powerful imaging techniques such as diffusion MRI (dMRI) can be used to map the structural connectivity between different brain regions [7–9]. At the macroscopic level, brain regions are delineated and defined as nodes of a network, while edges describe connectivity (structural or functional) between them. Consequently, the overall structure may be mathematically represented as a graph [2, 10]. Depending on the method used to construct the edges, brain networks can be divided into three types [2, 10, 11]: (i) structural networks that have edge weights based on strength of anatomical links between the nodes; (ii) functional networks in which the edges are given by statistical inter-dependence; (iii) functional networks whose edges are based on the causal influence of nodes. Clearly, the method employed to spatially parcellate the brain and consequently construct nodes will also affect the network parameters [12, 13]. The connectome has the potential to explain brain functions such as cognition, language, vision or audition, and their relation to specific brain regions since there exists a structure-to-function relationship between brain structural and functional networks [14].

In [15], ***robustness*** of the brain networks has been defined as the “degree to which the topological properties of a network are resilient to lesions such as the removal of nodes or edges.” In particular, robustness measures to what extent the brain can withstand damage from, or be affected by, lesions arising e.g. from tumor, trauma or stroke. Reduced robustness not only suggests potential for dysfunction due to the lesion, but may also indicate candidate target locations for treatment.

Brain resilience has been studied previously by considering the effect of deleting network nodes or edges from brain structural and functional networks, both computationally and empirically; see [16] for a comprehensive review. Brain robustness studies can broadly be divided into two categories. In the first, one attempts to predict lesion effects by computational models, i.e., virtually lesioning nodes and edges of the structural connectivity matrix and applying computational models to predict functional connectivity changes [17–19]. Subsequently, the predicted functional connectivity matrix can be compared with the empirical one and lesioning effects can be quantified using various graph measures. In the second category, one employs the empirical effects from brain lesions due to injury or disease. Studies using this approach focus on examining brain networks of patients with, e.g. traumatic brain injury (TBI), stroke or tumors, and quantify the effect of lesion location on the brain [20, 21], by comparison with data from age-and gender-matched healthy control subjects. Regardless of the approach, the structure-to-function network relation is utilized to predict the amount of damage which the brain can withstand due to lesions in a given location.

We apply the geometric notion of graph curvature to brain structural networks, and leverage this novel concept to analyze brain robustness. Previous studies have shown that network curvature can be used to differentiate cancer from normal tissue using gene co-expression networks [22], and to indicate market fragility in economic/financial networks [23]. It is important to note that, since network robustness can be viewed as the rate function at which a network returns to its original state after a perturbation, it has a positive correlation with entropy [24]. Consequently, network robustness and curvature are positively correlated through entropy [25].

### In this paper, we introduce the concept of graph curvature for studying brain structural connectivity networks

We use Ricci curvature and its contraction, the scalar curvature, on brain networks to introduce curvature as a nodal measure. In this study, using node curvature, we make two contributions to brain structural connectivity analysis: First, we identify areas of the brain that significantly contribute to the overall brain robustness, and hence we identify “important” nodes in brain networks. Previous studies have shown that hub nodes are critical for brain networks, but identifying such nodes is not straightforward. Node measures such as degree or strength do not identify all the hub nodes, and typically a combination of those measures, with centrality measures, is required [16, 26]. We show that node curvature not only corroborates findings based on strength and centrality measures, but additionally finds other key areas (e.g., inferior-frontal gyrus, middle-frontal gyrus, and inferior-temporal gyrus), which are not identified by any other node measure, and are important parts of the brain network. Second, by looking at the difference in node curvature, one can identify areas of the brain that are related to healthy development/aging, or abnormal neurodevelopment disorders such as autism spectrum disorders (ASD). In particular, we show that node curvature uniquely enables the identification of certain brain areas, with significantly affected structural connectivity in people with ASD.

The remainder of the paper is organized as follows. Section 2 provides some brief theoretical background on Ricci curvature, its relation to network robustness and curvature as a node measure for graphs. Details of the diffusion MRI data sets used in this study are also provided in this section. Section 3 presents results illustrating how brain curvature identifies: critical nodes (i.e. supporting robustness) in normal brain networks, nodes undergoing significant change in healthy development and aging, and nodes involved in abnormal development in ASD. Specifically, Section 3.1 shows critical nodes found using two different datasets of Diffusion Spectrum Imaging (DSI), which is a variant of dMRI, used by Hagmann et al. [26] to identify nodes constituting the structural core of the human cerebral cortex and by the MGH-USC HCP Consortium [9]. In Sections 3.3 and 3.4, we respectively use the WU-Minn HCP Consortium [7] Lifespan Pilot Project high angular resolution diffusion imaging (HARDI) data, and diffusion tensor imaging (DTI) data from the ASD study by Rudie et al. [27]. We describe changes in structural connectivity due to healthy development/aging, as well as ASD. We conclude by summarizing the results and future research directions in Section 4.

## 2. Methods

### 2.1 Wasserstein Distance and Optimal Mass Transport

Let *p* and *q* be two probability distributions on the discrete metric space *𝒳* equipped with metric *d*(,). The transportation cost of a unit mass from point *x*_*i*_ *∈ 𝒳* to *x*_*j*_ *∈ 𝒳* is denoted as *c*_*i,j*_ *≥*0. Denote by *π*_*i,j*_ *≥*0 the ***transference plan***, i.e., the (probability) measure of the amount of mass transferred from *x*_*i*_ to *x*_*j*_.

The Optimal Mass Transportation (OMT) problem seeks an optimal transference plan (*π*) that minimizes the total cost of moving *p* to *q*. This can be formulated as the following optimization problem [28–30]:

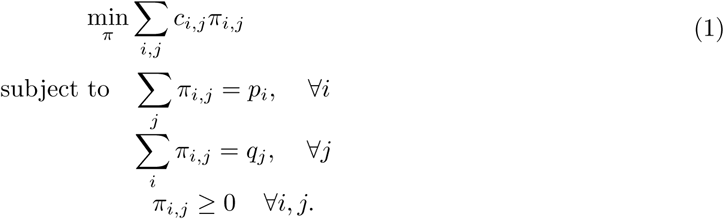

The problem in equation (1) may be expressed in the matrix form:

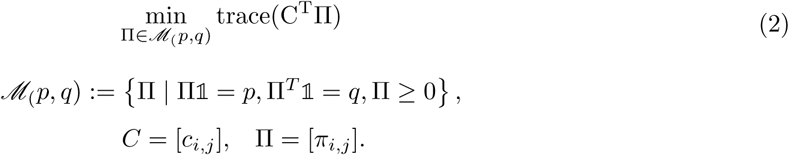

with

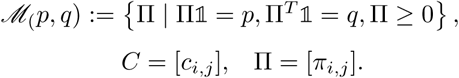

Here 1 is the column matrix of ones with the appropriate dimension.

When the cost *c* is defined as *c*_*i,j*_ = *d*(*x*_*i*_, *y*_*j*_)^*r*^, for any positive integer *r*, we can define *r*-Wasserstein distance [29, 31] as:

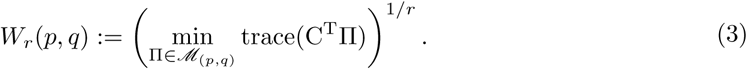

When *r* = 1 this is also known as the ***Earth Mover’s Distance (EMD).*** We will use this version of OMT in the present work.

### 2.2 Curvature

In this section, we introdcue the key notion of curvature from Riemannian geometry. Full details may be found in [32]. For *X* an *n*-dimensional Riemannian manifold, *x ∈ X,* let *T*_*x*_ denote the tangent space at *x*, and *v*_1_, *v*_2_ *∈ T_x_* orthonormal tangent vectors. Then, for geodesics (local curves of shortest length) *α*_*i*_(*t*): = exp(*tv*_*i*_), *i* = 1, 2, the sectional curvature *K*(*v*_1_, *v*_2_) measures the deviation of geodesics relative to Euclidean geometry, i.e.,

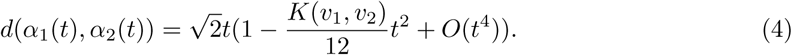

The Ricci curvature is the average sectional curvature. In other words, given a (unit) vector *u ∈ T_x_*, we complete an orthonormal basis *v, v*_2_, *…, v_n_*, and define 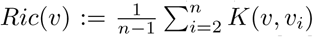. The Ricci curvature may be extended to a quadratic form known as the Ricci curvature tensor [32]. The scalar curvature is then defined to be the trace of this quadratic form.

There are a number of alternative characterizations of Ricci curvature [32]. In this paper, we employ the following definition: Referring to Figure 1, consider two very close points *x* and *y* in *X* and associated tangent vectors *w* and *′,* where *′* is obtained by parallel transport of *w* along a geodesic (in the direction *v*) connecting the two points. Now, geodesics drawn from *x, y* along *w, ′* will get closer when the curvature is positive (positively curved space). This is also reflected in the fact that the distance between two small (geodesic balls) is less than the distance of their centers. The Ricci curvature *Ric*(*v*) along the direction *v* connecting *x, y* quantifies this contraction. Similar considerations apply to negative and zero curvature.

**Figure 1:**
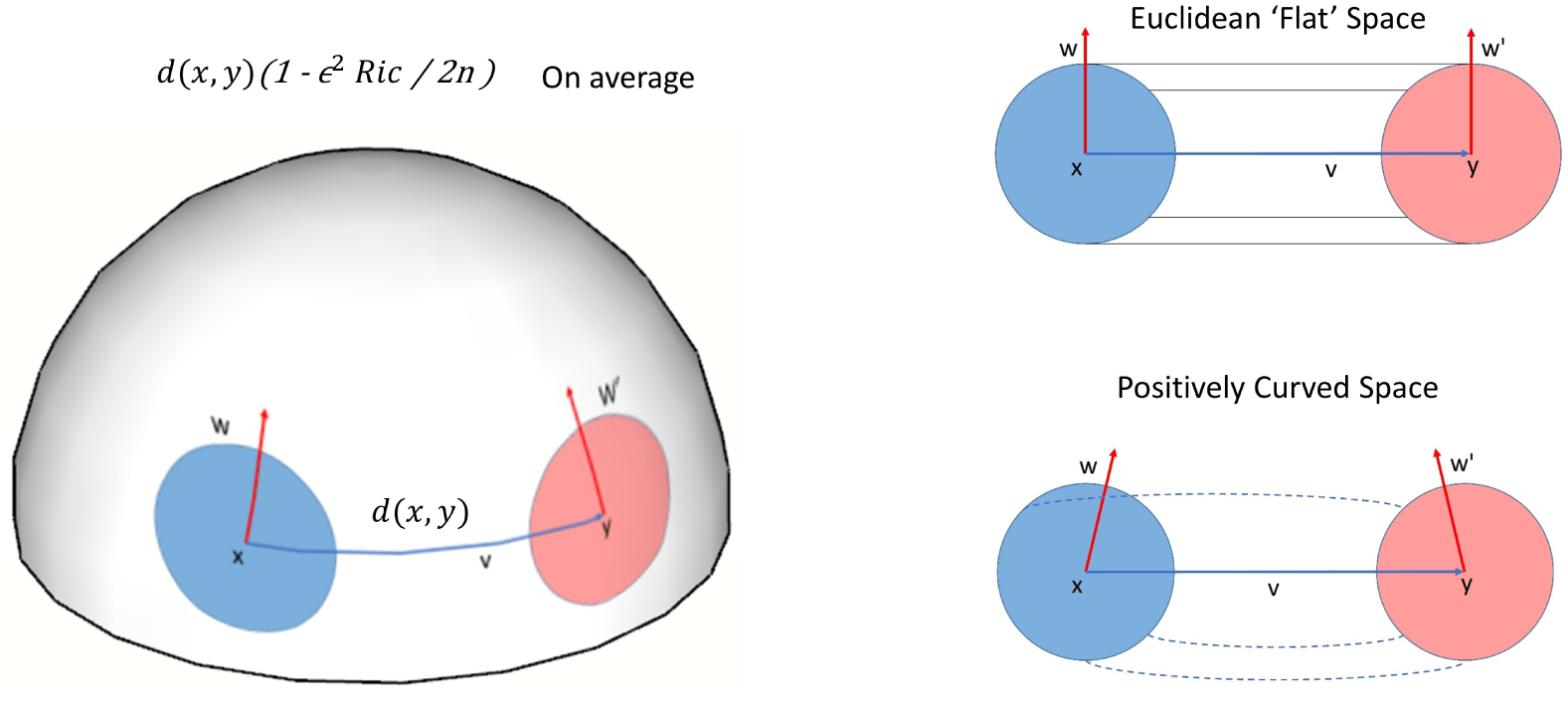
In a space with positive Ricci curvature, parallel geodesics emanating from points *x* and *y*, e.g., in directions along tangent vectors *w* (at *x*) and *w*^'^(at *y*), are drawn closer. In a Euclidean space, the distance between points moving along parallel geodesics at constant speed remains constant.

### 2.3 Ollivier-Ricci and Scalar Curvature

An analogue of the Ricci and of the scalar curvature exists for discrete spaces. Here, the place of the earlier geodesic (in a direction of a tangent vector *v*) connecting two points *x, y*, is taken simply by the pair (*x, y*). The so-called Ollivier-Ricci curvature is now denoted by *κ*(*x, y*) and the scalar curvature at *x* is the average *κ*(*x, y*) over neighboring nodes *y*. We now define these ideas in greater details.

#### 2.3.1 Ollivier-Ricci Curvature

***Ollivier-Ricci curvature*** or ***coarse Ricci curvature*** is the discrete generalization of the Ricci curvature [33–35]. Let (*X, d*) be a geodesic metric space equipped with a family of probability measures {*p*_*x*_: *x∈ X* }. We define the ***Ollivier-Ricci curvature*** *κ*(*x, y*) along the geodesic connecting *x* and *y* as

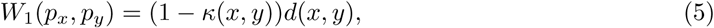

where *W*_1_ denotes the Earth Mover’s Distance (Wasserstein 1-metric), and *d* the geodesic distance on the space.

For the case of an undirected weighted graph (e.g. a brain structural connectivity network) *G* = (*V, E, W*), where *V* is the set of vertices (nodes), *E* the set of edges, and *W* the set of edge weights, we let

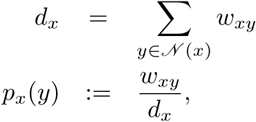

where *𝒩* (*x*) denotes the set of nodes that are adjacent to *x*; throughout we assume that all the edge weights *w*_*xy*_ = *w*_*yx*_ *≥* 0 and that *w*_*xy*_ = 0 *≥* if *d*(*x, y*) ≥ 2, or equivalently, if *y*∉ *𝒩* (*x*). Note here that the geodesic distance *d*(*x, y*) is taken to be the hop distance between node *x* and *y*, i.e., the minimum number of steps it takes to go from *x* to *y*.

#### 2.3.2 Node Curvature

The ***(scalar) node curvature*** for node *x* (*κ*_*x*_) in the graph is computed by summing the curvature between node *x* and all its neighboring nodes, i.e.,

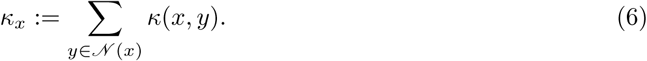

We also note that an alternative “weighted” version of the node curvature may be defined as

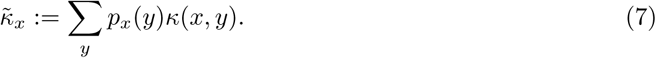

### 2.4 Robustness and the Fluctuation Theorem

We now turn to the notion of robustness which we will employ in this paper, and subsequently make the link between robustness and curvature. It is based on ideas from statistical mechanics and, in particular, the ***Fluctuation Theorem*** formulated in [24]. The Fluctuation Theorem measures the ability of a network to maintain its functionality in the face of perturbations (internal or external).

Let *p*_*Δ*_(*t*) be the probability that the mean value of an observation (for a given network) deviates from its original value, by more that *Δ* at time *t*, due to some perturbation. The ***rate*** *R* at which the system returns back to its original state is defined as

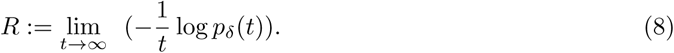

Note that large *R* means not much deviation and small *R* implies a large deviation. In statistical mechanics, it is well-known that entropy and rate functions from large deviations are very closely related [24,36]. The Fluctuation Theorem is a mathematical statement relating the positive correlation of changes in system entropy Δ*S* to changes in robustness Δ*R*:

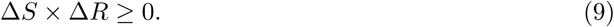

### 2.5 Ollivier-Ricci Curvature and Graph Robustness

In this section, following [22, 23], we establish the relationship between curvature and robustness. For ease of explanation, we consider *X* to be a Riemannian manifold; the relations drawn can be extended to discrete spaces.

It turns out that one can endow the space of probability densities on *X* (taken with respect to the volume measure) with a natural Riemannian structure as well; see [37] for details. We denote the space of probability densities by *𝓅* (*X*). Then, one defines the ***Boltzmann entropy*** of *ρ ∈ 𝓅* (*X*) as

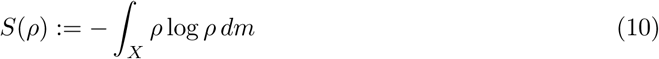

where *dm* denotes the volume measure on *X*. (There are several technical assumptions that should be made to ensure the existence of *S*, see [25, 38].)

We can then express the following remarkable result from [25, 38]: Let *Ric* denote the Ricci curvature defined on *X*, and suppose the *Ric≥ k* for any tangent vector on *X*. Then, for every *ρ*_0_, *ρ*_1_ *∈ 𝓅* (*X*), the constant speed geodesic *ρ*_*t*_ with respect to the Wasserstein 2-metric connecting *ρ*_0_ and *ρ*_1_ satifsfies

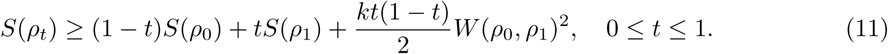

This relationship indicates the positive correlation of of changes in entropy and curvature [22], namely

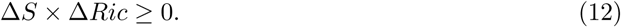

Similar arguments can be made on discrete spaces. Following the arguments of [22, 23], and the Fluctuation Theorem, one can express the positive correlation of graph Ricci curvature (see equation (5)) and robustness, expressed as:

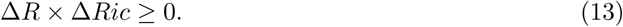

***The main conclusion is that one can use network curvature as a measure of functional robustness***. Since curvature can easily be computed via linear programming [28, 30], it provides an attractive and novel tool to characterize the robustness of networks represented as weighted graphs, such as brain connectivity networks. In the next section, we briefly summarize existing measures to characterize complex brain networks and provide information about the datasets which will be used next to demonstrate the benefits of curvature.

### 2.6 Measures of Brain Networks Characteristics

We hereafter briefly summarize important graph-theoretical measures, which have been introduced to characterize brain networks [39], and will be used in our experiments.

#### 2.6.1 Node Strength *s*_*i*_

The strength of a node *i* is the sum of the weights *w*_*ij*_ of the node’s adjacent edges [40], i.e.,

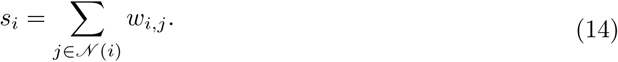

Then dMRI data may be employed, in combination with deterministic or probabilistic integration/propagation methods of vector fields (called *tractography*), to assess the likelihood of connections between cortical and sub-cortical areas [41]. Such likelihood can be obtained by the number of three-dimensional curves generated by these integration or propagation methods and used, in the context of brain structural networks, to define the weight *w*_*i,j*_ of an edge between two nodes *i* and *j*. This summarizes how strongly connected those nodes are to each others, and to the rest of the brain.

### 2.6.2 Betweenness Centrality *g*_*i*_

The betweenness centrality of a given node *i* is defined as the number of shortest paths between pairs of nodes that pass through the node *i* [42], i.e.,

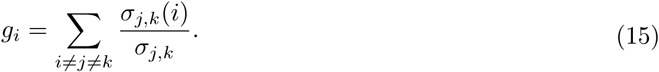

Where *σ*_*j,k*_ is the total number of shortest paths from node *j* to node *k* and *σ*_*j,k*_(*i*) is the number of those paths that pass through *i.*

#### 2.6.3 Clustering Coefficient *C*_*i*_

The Clustering Coefficient of node *i* is a measure of the density of connections among the node’s topological neighbors [43, 44]. This is defined as follows: Take *i ∈V*, the vertex set of a graph *G* = (*V, E, W*), and assume unit weights *e*_*ij*_ *∈W* for all existing edges. Suppose that the node *i* has *k*_*i*_ neighbors. For an un-directed graph (which is usually the case for brain structural connectivity networks), there can be at the most *k*_*i*_(*k*_*i*_ -1)*/*2 edges in the local neighborhood. Then, *C*_*i*_ is defined as the fraction of the edges that actually exist between the immediate neighbors of *i* over the maximal number of edges, i.e.,

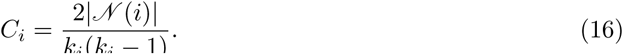

As before, *𝒩* (*i*) = {*j*: *e*_*ij*_ *∈E*} is the immediate neighborhood of nodes if *i*, and |*𝒩*| denotes the cardinality of this set.

### 2.7 Diffusion MRI Datasets

As briefly described in the introduction, we used four different public/open access datasets in our experiments, and we now provide more details about this data. First, we assess the ability of node curvature to capture novel information, which is complementary to existing measures of brain networks characteristics (e.g. strength). We analyze the high-resolution connectivity matrices created and analyzed by Hagmann et al [26], using DSI data from 5 healthy subjects. These matrices are available from the USC Multimodal Connectivity Database [45], which enables the reproduction of the original results [26], and evaluation of our method with the exact same datasets. We also analyze 33 new DSI datasets obtained from the MGH-USC HCP Consortium [9], to demonstrate the consistency of our findings on critical brain areas. Next, we illustrate the ability of node curvature to capture changes in certain brain areas, which are related to healthy development and aging. These experiments use HARDI datasets obtained from the WU-Minn HCP Consortium Lifespan Pilot Project [7]. Finally, we show that node curvature is uniquely capable of detecting changes in brain structural connectivity in ASD, which are in agreement with previous morphometric MRI studies. These last experiments are performed with DTI datasets from 51 subjects with ASD and 43 typically developing subjects, as previously published by Rudie et al. [27]. The connectivity matrices from this study are also available from the USC Multimodal Connectivity Database [45], which again enabled straightforward reproduction of those results and comparison with our method.

#### 2.7.1 DSI Datasets from Hagmann et al. [26]

Data were acquired in 5 healthy right-handed male subjects, on a Philips Achieva 3T scanner with voxel size 2 × 2 × 3 *mm*3, TR/TE=4200/89 ms and 129 diffusion gradients with a maximum *b*-value of 9000 *s/mm*2, for a total acquisition time of 18 min. After segmentation of the white and gray matter, 998 cortical regions-of-interest were created, with an average size of 1.5 *cm*2. Tractography was then performed, and structural connectivity matrices created by defining the weight of each edge as the number of streamlines per unit surface (i.e. density). Additional details can be found in the original paper [26].

#### 2.7.2 DSI Datasets from the MGH-USC HCP Consortium

Data were acquired in 35 healthy adults (age range 20 to 59) scanned on the customized Siemens 3T Connectom scanner and are available at https://db.humanconnectome.org. Two of the datasets were not included in our experiments because of pre-processing errors in our analysis pipeline. Acquisition parameters included voxel size of 1.5 1.5 1.5 *mm*3, TR/TE=8800/57 ms and four *b*-values (with corresponding number of diffusion gradients in parenthesis): 1000 *s/mm*2 (64), 3000 *s/mm*2 (64), 5000 *s/mm*2 (128), 10000 *s/mm*2 (256), for a total acquisition time of about 89 min. Connectivity matrices were generated using DSI Studio (http://dsi-studio.labsolver.org) as described below.

#### 2.7.3 HARDI Datasets from the WU-Minn HCP Consortium Lifespan Pilot Project

Data were acquired from healthy subjects across the lifespan in 6 age groups: 4-6, 8-9, 14-15, 25-35, 45-55 and 65-75 years and are available at https://db.humanconnectome.org. We analyzed the data acquired on the UMinn Siemens 3T Prisma scanner (Phase 1a), which include 5 participants per age group (ages 25-35, 45-55 and 65-75) or 6 participants per age group (ages 8-9 and 14-15). Acquisition parameters included voxel size of 1.5 × 1.5 × 1.5 *mm*3, TR/TE=3222/89 ms and two *b*-values, 1000 *s/mm*2 and 2500 *s/mm*2, each with 75 diffusion gradients acquired twice with opposite phase-encoding polarity, for a total acquisition time of about 22 min. Connectivity matrices were also generated using DSI Studio (http://dsi-studio.labsolver.org) as described below.

#### 2.7.4 DTI Datasets from Rudie et al. [27]

Data were acquired in 51 subjects with ASD and 43 typically developing subjects, on a Siemens 3T Trio scanner with voxel size 2 × 2 × 2 *mm*3, TR/TE=9500/87 ms and one *b*-value of 1000 *s/mm*2 with 32 diffusion gradients, for a total acquisition time of about 8 min. Deterministic tractography was used to create structural connectivity matrices, with weights defined as the number of streamlines connecting each pair of 264 cortical areas (nodes). Additional details can be found in the original paper [27].

#### 2.7.5 Generation of Connectivity Matrices for the HCP Datasets

We used DSI Studio (http://dsi-studio.labsolver.org) [46] to process the HCP DSI and HARDI datasets.

To run tractography and generate connectivity matrices for the DSI data, seeds were placed randomly in the whole brain with the following settings: normalized quantitative anisotropy (NQA) threshold: 0.08, angular threshold: 60^*m*^, tractography method: Runge-Kutta [47], total number of streamlines: 5 million (Although similar results were obtained with 500, 000 streamlines, we used 5 millions to ensure consistency with [26]). 116 cortical areas (nodes) were automatically segmented via non-linear registration of the Automated Anatomical Labeling (AAL) template available in DSI Studio.

For the HARDI data, diffusion tensors were estimated to perform tractography. Seeds were also placed randomly in the whole brain with the following settings: fractional anisotropy (FA) threshold: 0.1, angular threshold: 60° , tractography method: Runge-Kutta [47], total number of streamlines: 500, 000. 116 cortical areas (nodes) were automatically segmented via non-linear registration of the Automated Anatomical Labeling (AAL) template available in DSI Studio.

## 3. Results

### 3.1 Curvature as a Hallmark of Brain Areas Robustness

Individual node curvature (as defined in Section 2.3.2) of brain regions contributes to the overall (average) curvature of the brain network. This measure not only helps identify changes in the network, as discussed in the previous sections, but also can help identify relatively more important parts of the brain structural network. As explained in Section 2.5, equation (8), curvature is directly correlated to network robustness. Therefore, nodes with higher curvature contribute more to the overall structural robustness of the network.

To demonstrate this, we performed experiments using two different Diffusion Spectrum Imaging datasets: First, the DSI data for 5 participants, as presented in [17, 26], was considered, to enable comparison of our results to previous studies. High-resolution connectivity matrices (998 × 998) were obtained from the USC Multimodal Connectivity Database [45]. Second, the DSI data for 33 participants from the MGH-USC HCP Consortium was also employed, and lower resolution connectivity matrices (116 × 116) were generated as described in Section 2.7.2.

As previously described, we emphasize here that comparing properties across brain networks with different resolutions (i.e. number of nodes) should be done only with great care [12], as brain network properties can differ significantly with nodal parcellations [12, 13]. Nonetheless, it is worth studying curvature as a measure which may provide information across different network resolutions: High resolution parcellations, also known as *dense connectomes* [48] will ultimately provide greater insights into the structure of brain networks, while lower resolution parcellations are more easily manageable, since they requires less computational resources.

First, we present results based on the high resolution connectivity matrices [45] in Figure 2. Here, we show the top 25% of nodes with the highest node curvature, strength and betweenness centrality, appearing consistently across 5 participants. Figures in panels B and C follow the same convention as Figure 2 (C) and Figure 7 (A) of [26] respectively, and are presented here for comparison purposes. The details of the areas identified by all three measures can be found in Supplementary Note 1. In previous study [26], using several network analysis methods, eight anatomical sub-regions were identified as belonging to the so-called *structural core network* of the human brain. These include the posterior cingulate cortex, precuneus, cuneus, paracentral lobule, isthmus of the cingulate, banks of the superior temporal sulcus, and inferior/superior parietal cortex, all of them in both hemispheres. Additionally, [17] showed that lesions in the temporo-parietal junction, cortical midline and frontal cortex have the most extensive effects on brain functionality. Also, we note that the medial prefrontal cortex forms part of the default mode network of the human brain [49]. Panel A of Figure 2 shows that curvature identifies areas in the inferior-frontal gyrus, middle-frontal gyrus and inferior-temporal gyrus, consistent with [26] [17], and thus providing very interesting information based on network structure, which is not captured by strength or betweenness centrality.

**Figure 2:**
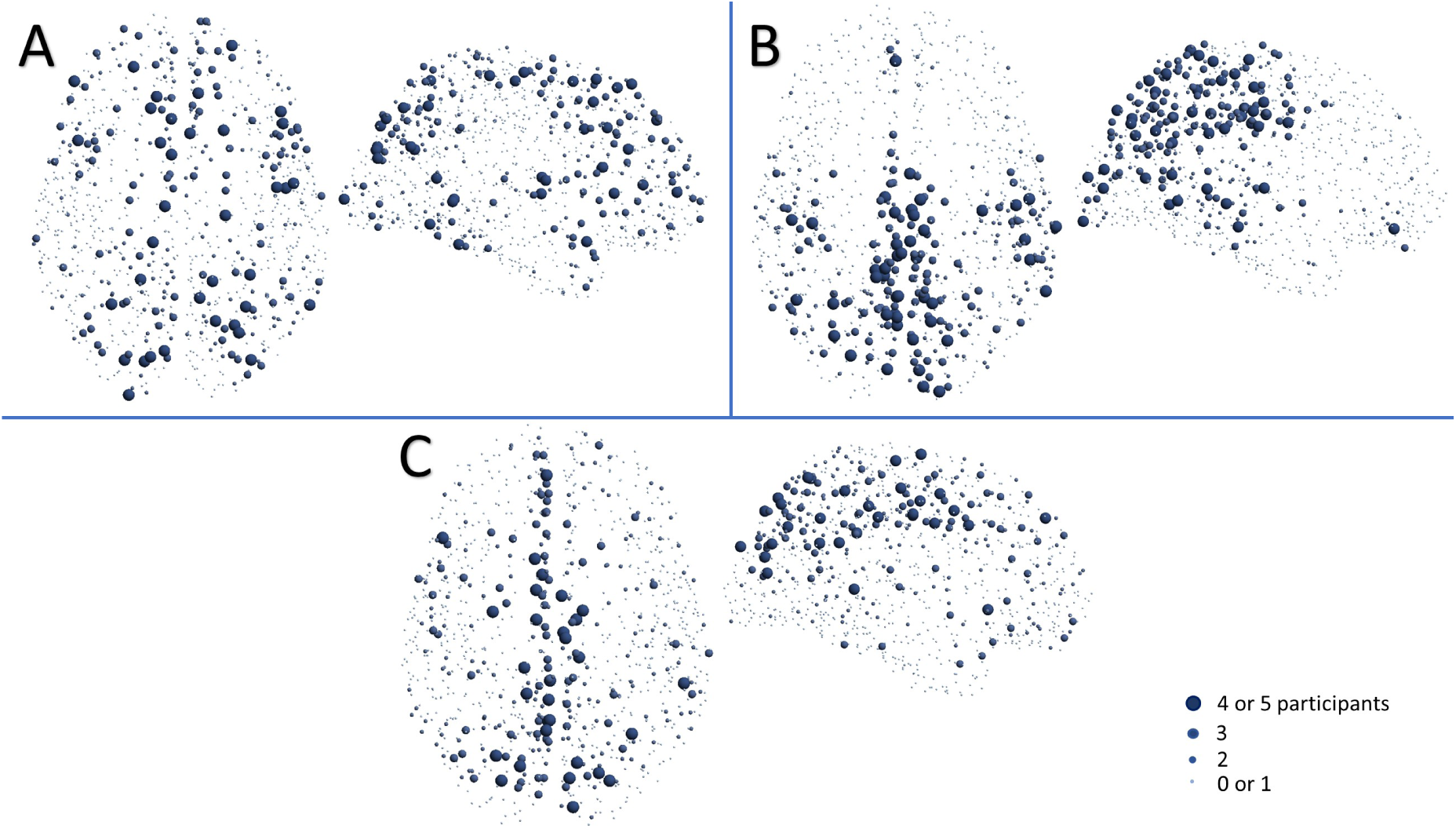
Nodes (top 25%) with highest curvature (A), strength (B) or betweenness centrality (C) consistently present across subjects. For instance, the largest spheres indicate nodes with high values in 4 or 5 out of the 5 subjects. Results from high resolution connectivity matrices (998 × 998) from Hagmann et al [26].

Second, following the same organization as Figure 2, Figure 3 shows results for the low resolution matrices generated from the MGH-USC HCP Consortium datasets. As expected, distinct areas are identified with all three measures (since cortical parcellation is different from the one used in Figure 2) [12]. We should also note that the high resolution data did not include the cerebellum. Nodes with high strength and betweenness centrality are found more towards the frontal, precentral, superior parietal areas and in the cerebellum. Once again, curvature supplements the information provided by other measures and identify areas in the inferior-frontal gyrus and transverse temporal gyrus (Heschl’s gyrus) in both hemispheres, where lesions are known to induce pronounced effects in loss of brain functionality [17] (see the list of areas in Supplementary Note 2).

**Figure 3:**
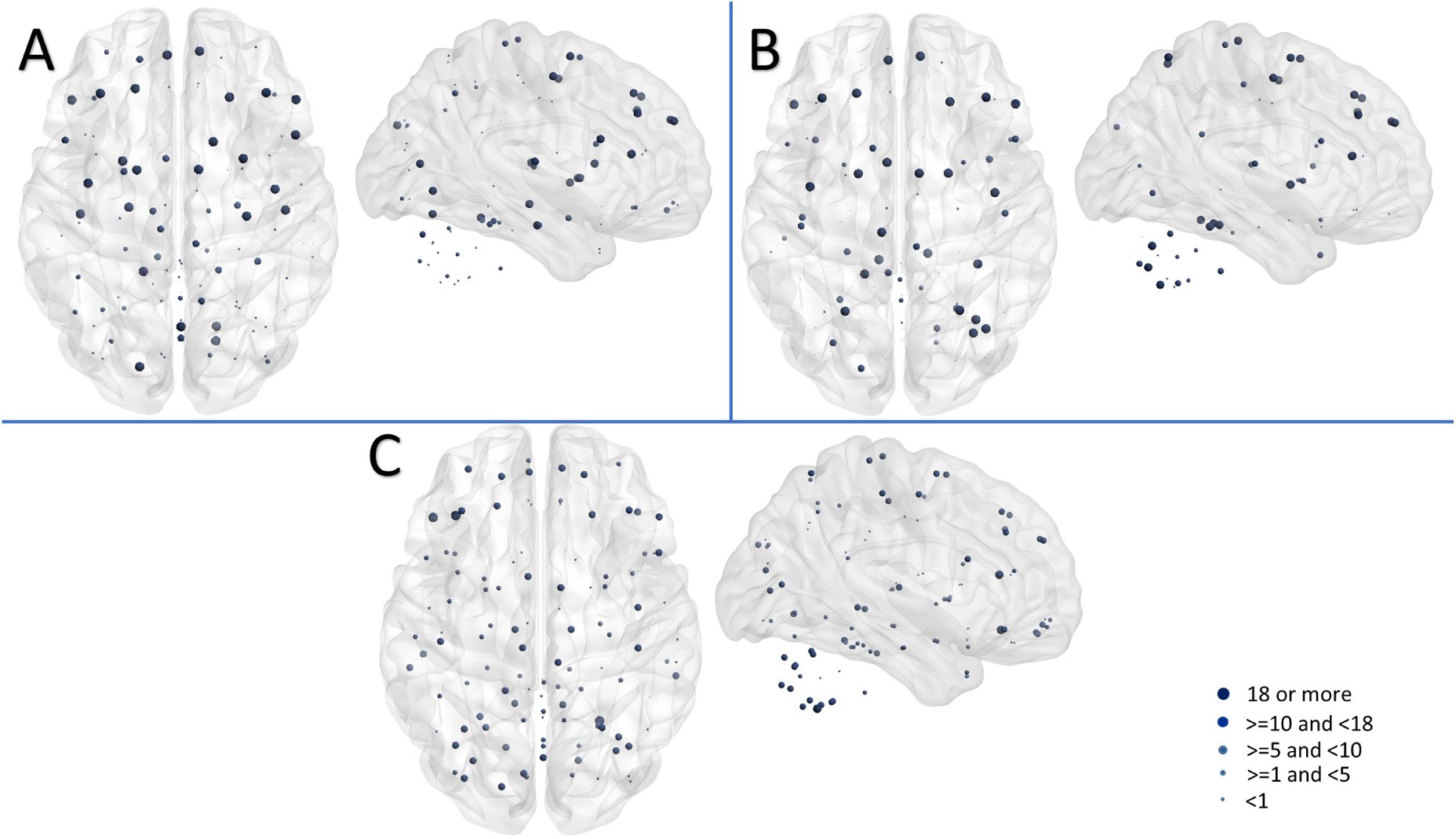
Nodes (top 25%) with highest curvature (A), strength (B) or betweenness centrality (C) consistently present across subjects. For instance, the largest spheres indicate nodes with high values in 18 out of the 33 subjects. Results from lower resolution connectivity matrices (116 × 116) generated using the AAL atlas and MGH-USC DSI datasets.

### 3.2 Remarks on Graph Measures and Assessment of Brain Network Robustness

In order to quantify the importance of a given node in a brain network, the effect of node(s) deletion on graph measures can be considered [17]. Based on node measures such as strength, betweenness centrality, and curvature, given node(s) and all related edges are removed (i.e., by deleting the row and the column) from the connectivity matrix. Independent graph measures such as connectedness or global efficiency are then computed on the remaining connectivity matrix, and the effect of node(s) and edges removal is quantified by the faster or slower decay of those measures. For instance, if a network becomes disconnected (i.e. size of largest connected component becomes small) quickly, when removing nodes based on their importance, as quantified by a particular graph measure, we can conclude that this graph measure provides an effective way to identify important nodes (because the graph breaks down quickly after removing only a few nodes). If, however, the graph becomes disconnected slowly when removing nodes based on a different graph measures, we can conclude that this particular is not as good as identifying important nodes.

#### 3.2.1 Gaussian Transformation of the Connectivity Matrix Weights

Streamlines numbers (i.e., the weights of structural connectivity graphs) produced by tractography algorithms are exponentially distributed [12, 26]. Without altering the rank-ordering of strong to weak pathways, those distributions can be transformed to Gaussian distributions [17] for ease of analysis. However, we would like to emphasize that such transformation may lead to changes in edge weights (and consequently node strengths) that may affect graph measures differently, as illustrated next and thereby possibly biasing the definition of important nodes, as well as the identification of graph measures particularly adequate to quantify this importance. For instance, the weights, while preserving order, can be mapped to a completely different range of values, thereby increasing or diminishing the relative importance of nodes. (e.g. mapping weights 1 and 1000 to 10 and 11). We therefore recommend to apply such transformation(s) with care.

Using the size of largest graph component and global efficiency as metrics (i.e., the independent graph measures), as in [17] (see e.g., Figure 3 in this paper), Figures 4 and 5 illustrate the fact that if such a Gaussian transformation is performed [17], betweenness centrality can be seen as a better measure of node importance compared to strength (see Figures 4 and 5, panels C-D). Both the largest graph component and betweenness centrality decrease rapidly. However, without this transformation (i.e, with original connectivity matrices), although the largest graph component preserves its behavior with deletion of nodes based on either strength or betweenness centrality, global efficiency does not: strength appears to be a better measure of node importance (see Figures 4 and 5, panels A-B). We note that, despite having very different numbers of nodes, both DSI Datasets from Hagmann et al. [26] and the MGH-USC HCP Consortium exhibit the same results. This illustrates the potential effect of transforming connectivity matrix weights, and provides justification for our choice to generally **not** transform those weights.

**Figure 4:**
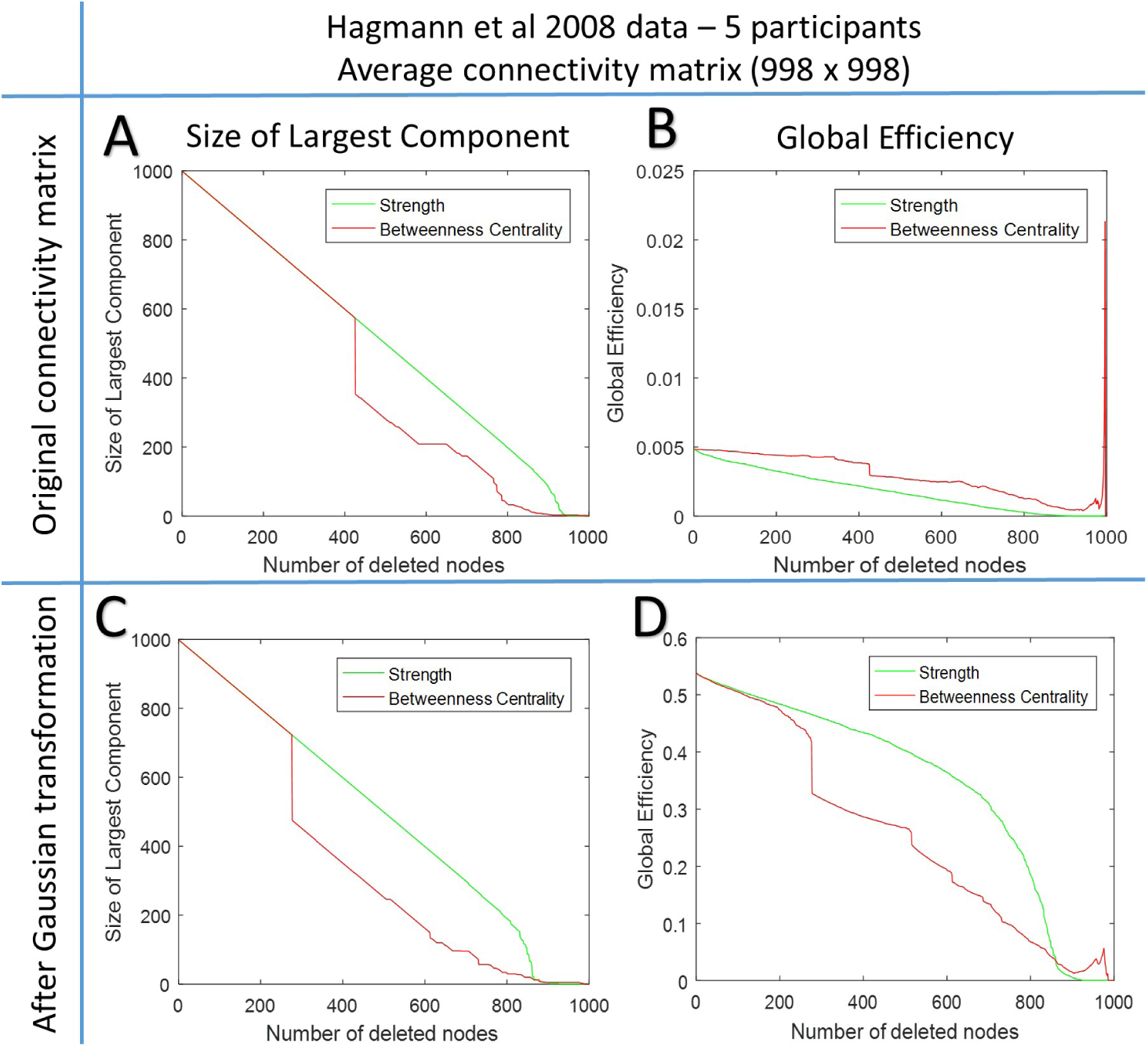
Robustness analysis using node deletion based on their strength or betweenness centrality, for the high resolution connectivity matrices (998 × 998) from Hagmann et al [26]. The size of the largest component and global efficiency are computed (with or without transformation of the connectivity matrix weights) after targeted removal of nodes with high strength or betweenness centrality. The top row shows results for the original connectivity matrix while the bottom row shows results after Gaussian transformation of its weights. Note that results shown in panel C and D are similar to Fig. 3 in [17].

**Figure 5:**
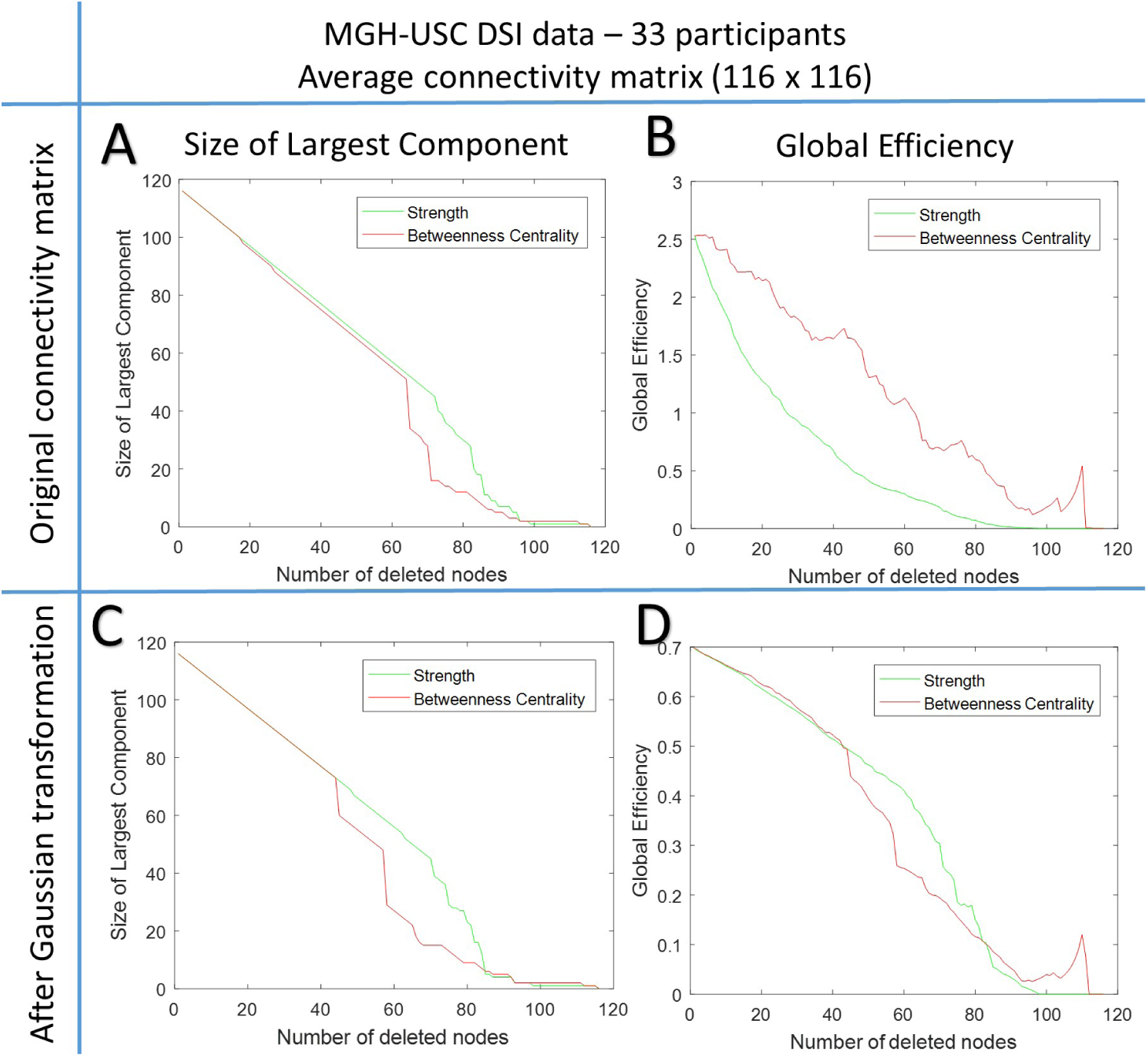
Robustness analysis using node deletion based on their strength or betweenness centrality, for the lower resolution connectivity matrices (116 × 116) generated using the AAL atlas and MGH-USC DSI datasets. The size of the largest component and global efficiency are computed (with or without transformation of the connectivity matrix weights) after targeted removal of nodes with high strength or betweenness centrality. The top row shows results for the original connectivity matrix while the bottom row shows results after Gaussian transformation of its weights. Note that results are consistent with those presented in Figure 4.

#### 3.2.2 Comparison Across Nodal Scales

For each step of a robustness analysis as presented in Figures 4 and 5, a node is deleted and the size of the network is reduced. Thus, we end up comparing parameters across brain networks with different parcellations (i.e. different number of nodes). It is important to note that network topological measures show strong dependence on the nodal scale [12], therefore, discrepancy in the measures may be attributed, to some extent, to disparate nodal scales.

#### 3.2.3 Graph Measures to Assess Node Robustness

Traditionally, integration and centrality graph metrics like global efficiency, betweenness centrality, degree centrality, characteristic path length, and clustering coefficient have been used to study the robustness of brain networks [16]. While these measures certainly provide very useful information about the networks, we argue, based on the discussion provided in Section 2.5, that graph curvature and entropy may be more relevant metrics for robustness. To explore this, Figure 6 displays the changes in topological entropy^1^ when nodes are deleted based on **decreasing** strength, curvature and betweenness centrality. For this experiment, the average of the 5 MGH-USC HCP Consortium DSI connectivity matrices (as described in Section 2.7.2) was used. Nodes with least importance, i.e., lower curvature, strength and betweenness centrality were deleted first, therefore node measures maintaining the entropy after the most number of nodes removal should give a better indication of robustness. We observe that node curvature and strength show similar results, as entropy remains high as more nodes (with low strength or curvature) are deleted, as compared to betweenness centrality. Thus in both cases, i.e., the original connectivity matrix (Figure 6, panel A) and the matrix after the Gaussian transformation (Figure 6, panel B), curvature and strength identify nodes contributing more towards the overall graph robustness.

**Figure 6:**
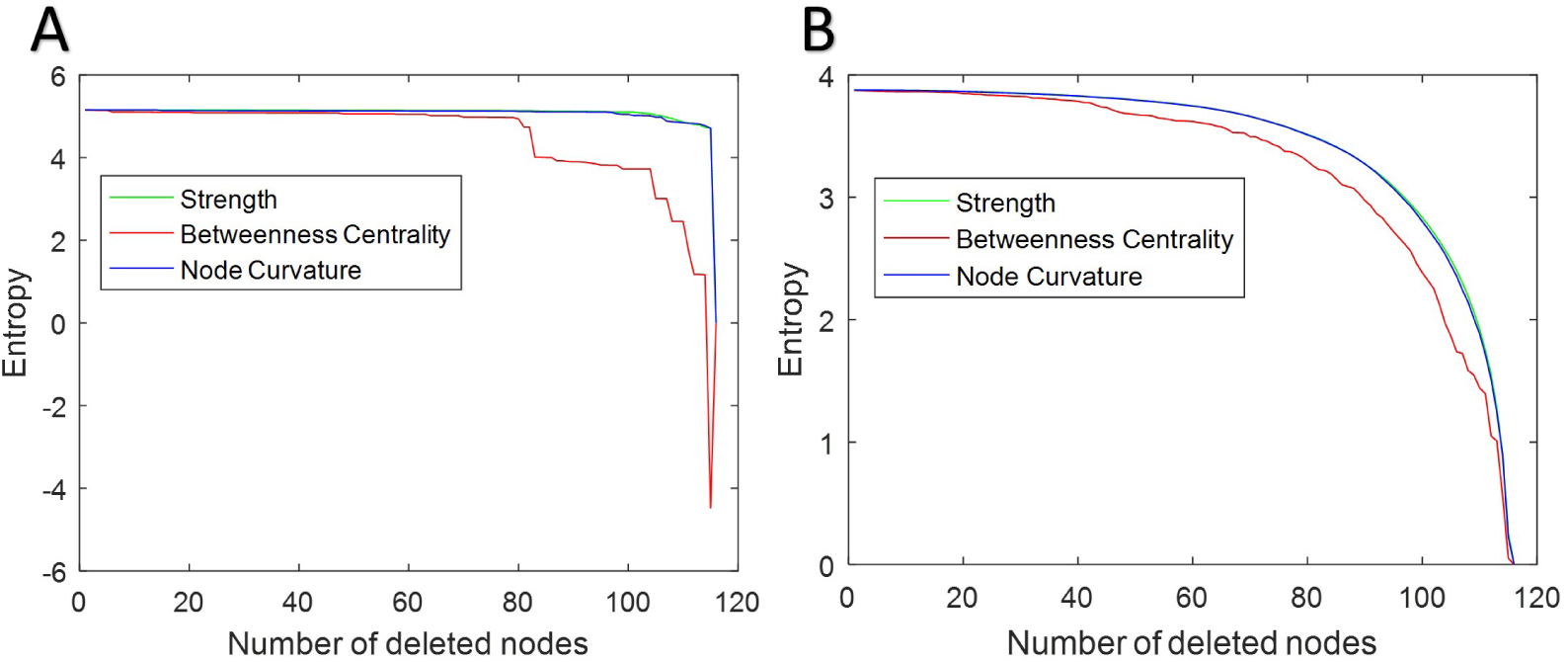
Robustness analysis on the basis of node deletion with topological entropy as a measure, using low resolution average connectivity matrix from MGH-USC DSI data. (A) Original average connectivity matrix (B) After Gaussian transformation of the connectivity matrix. Note that Node Curvature and Strength show similar trend and are better measures of robustness as compared to Betweenness Centrality.

**Figure 7:**
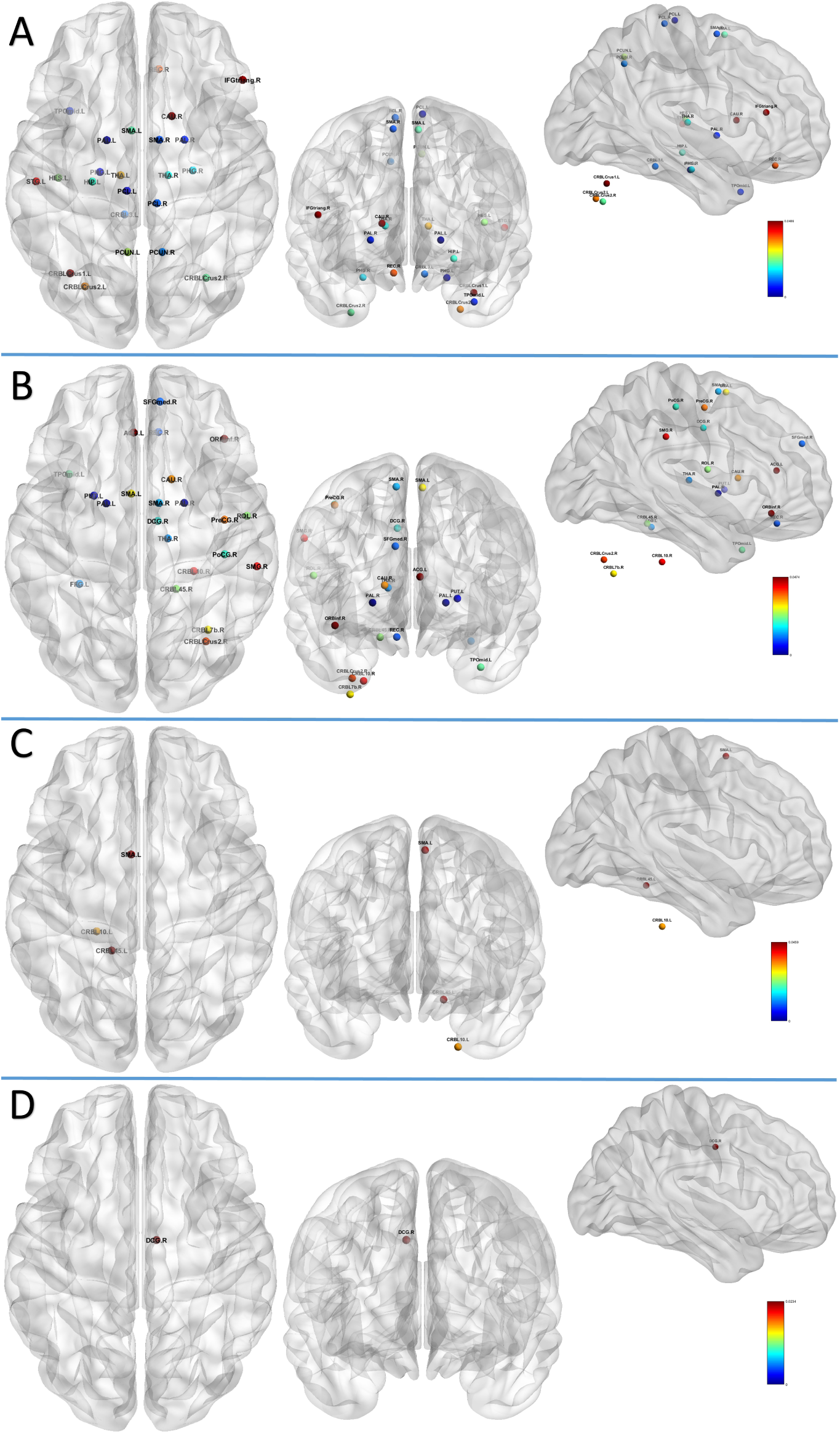
Nodes with significant difference in structural connectivity related to healthy aging. Node color correspond to the *p*-value. (A) Node curvature. (B) Node strength. (C) Betweenness centrality. (D) Clustering coefficient. **Node Name Abbreviations**: Suffix R = Right hemisphere, L = Left hemisphere; ACG = Anterior cingulate gyrus; CAU = Caudate nucleus; CRBL = Cerebellum; CRBLCrus = Crus of cerebellar hemisphere; DCG = Mid-cingulate area; FFG = Fusiform gyrus; HES = Heschl’s gyrus; HIP = Hippocampus; IFGtriang = Pars triangularis; ORBinf = Pars orbitalis; PAL = Pallidum; PCL = Paracentral lobule; PCUN = Precuneus; PHG = Parahippocampus; PoCG = Postcentral gyrus; PreCG= Precentral gyrus; PUT = Putamen; REC = Rectus gyrus; ROL = Rolandic operculum; SFGmed = Medial frontal gyrus; SMA = Supplementary motor area; SMG = Supramarginal gyrus; STG = Superior temporal gyrus; THA = Thalamus; TPOmid = Middle temporal pole.

### 3.3 Change in Curvature and Healthy Development/Aging of the Brain

We used datasets from the WU-Minn HCP Consortium Lifespan Pilot Project to study structural changes in brain networks related to healthy aging. Details about the data and construction of connectivity matrices are given in Section 2.7.3. Although node measures such as strength, betweenness centrality, and clustering coefficient can provide useful information about areas involved in healthy aging, we focus mainly on information provided by curvature in the present discussion. In Figure 7, we summarize the most pronounced changes related to healthy aging, and show results for the differences in node measures (see the list of areas in Supplementary Note 3) between the Lifespan group 1 (age 8-9) and group 5 (age 65-75). Figure 7 (panel A) illustrates that curvature finds significant bilateral differences in the parahippocampal gyrus, precuneus, paracentral lobule, thalamus and cerebellum. Additionally, curvature can detect differences in the hippocampus, Heschl’s gyrus, and temporal areas in the left hemisphere, as well as the caudate nucleus, gyrus rectus, frontal inferior gyrus, pars triangularis in the right hemisphere. Those findings are in agreement with previous studies demonstrating age related structural changes in the thalamus and hippocampus [51,52], cerebellum [52], precuneus [53], para-hippocampus and precentral gyrus [54].

**Exploratory statistical analysis**: We note that in order to evaluate the statistical significance of differences for node measures (e.g. curvature) between two groups (e.g., different age groups, or controls and patients below), which can be of different sizes, unpaired two-sample *t*-tests were conducted. The null hypothesis of equal means between groups, and for each node, was tested and rejected for *p*-values (uncorrected for our exploratory results) less than 0.05, without assuming equal variances.

### 3.4 Change in Curvature and Autism Spectrum Disorders (ASD)

The aim of this experiment is to test whether various measures of node importance/robustness (curvature, strength, centrality and clustering) can detect differences in structural connectivity between individuals with Autism Spectrum Disorders (ASD) and typically developing subjects (TD). We utilized DTI data from 51 individuals with ASD and 43 TD participants, from a previously published study [27], so as to compare ASD-related differences in the brain network organization. Details about the data are given in Section 2.7.4. The DTI connectivity matrices capturing the brain structural connectivity of each participant were generated using a set of 264 coordinates [55] in MNI space. Additional details about the scanning protocol, diffusion data pre-processing and generation of connectivity matrices may be found in [27].

The original study [27] identified both functional and structural networks differences related to autism. For structural networks, changes in the white matter integrity and streamlines count were noted. Further, age-related atypical changes in the balance of local and global efficiency in structural and functional networks were observed. However, six global graph-theoretic metrics, namely the clustering coefficient, normalized clustering coefficient (*λ*), characteristic path length (CPL), normalized CPL (*γ*), small-worldness and modularity did not show statistically significant differences between subject the ASD and TD groups [27]. Interestingly, we found that strength, betweenness centrality, clustering coefficient and node curvature are able to identify areas with significant differences between groups. Therefore, we focus in these last four measures and summarize our findings next.

Figure 8 shows nodes with statistically significant (see remark above) differences between the ASD and TD groups, using node measures: curvature, strength, betweenness centrality and clus-tering coefficient (see the list of areas in Supplementary Note 4). While node strength, betweenness centrality and clustering coefficient provide important information about affected areas, we should note that **curvature identify differences in areas that are not detected by the other measures**. In particular, curvature discovered differences in the following areas of the right hemisphere: inferior temporal gyrus (temporo-occipital), temporal occipital fusiform cortex, para-hippocampal gyrus (posterior), cuneus cortex, supra-marginal gyrus (posterior), insular cortex, frontal orbital cortex, and frontal pole (Figure 8, Panel A), which is in line with previous studies [56–60]. Interestingly, a morphometric analysis of asymmetry in volume/structure of cortical areas in individuals with ASD found a pronounced rightward bias [56], which appears to agree with our findings based on curvature. Indeed, node curvature detected all the affected areas (in case of both ASD and developmental language disorder (DLD)) in the right hemisphere, except for the lingual gyrus, as also reported in Table 4 of [56].

**Figure 8:**
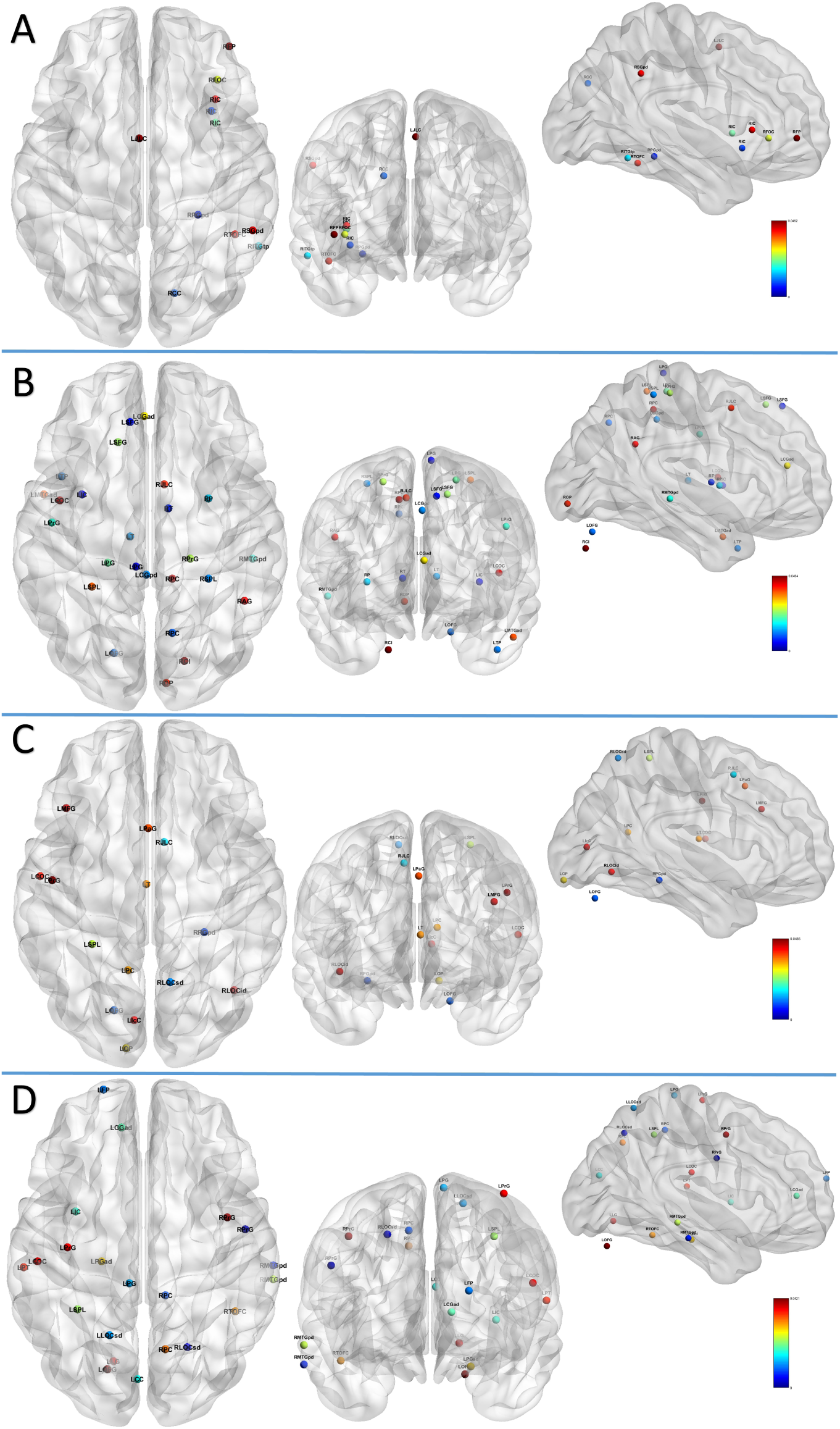
Nodes with significant difference in structural connectivity between individuals with Autism Spectrum Disorders and typically developing subjects. Node color correspond to the *p*-value. (A) Node curvature. (B) Node strength. (C) Betweenness centrality. (D) Clustering coefficient. **Node Name Abbreviations**: Suffix R = Right hemisphere, L = Left hemisphere; AG = Angular gyrus; CC = Cuneus cortex; CGad = Cingulate gyrus, anterior division; CGpd = Left cingulate gyrus, posterior division; COC= Central opercular cortex; FOC = Frontal orbital cortex; FP = Frontal pole; IC = Insular cortex; IcC = Intracalcarine cortex; ITGtp = Inferior temporal gyrus, temporo-occipital part; JLC = Juxtapositional lobule cortex; LG = Lingual gyrus; LOCid = Lateral occipital cortex, inferior division; LOCsd = Lateral occipital cortex, superior division; MTGad = Middle temporal gyrus, anterior division; MTGpd = Middle temporal gyrus, posterior division; OFG = Occipital fusiform gyrus; OP = Occipital pole; P = Putamen; PaG = Paracingulate gyrus; PC = Precuneus cortex; PG = Postcentral gyrus; PGad = Parahippocampal gyrus, anterior division; PGpd = Parahippocampal gyrus, posterior division; PrG = Precentral gyrus; SFG = Superior frontal gyrus; SGpd = Supramarginal gyrus, posterior division; SPL = Superior parietal lobule; T = Thalamus; TOF = Temporal occipital fusiform cortex.

**4.Conclusion**

In this paper, we have introduced the concept of graph curvature to quantify the importance of nodes (meaning that their disruption leads to large changes in the overall graph) in brain networks. We have shown that curvature can indeed help identify important areas/nodes and points to changes in the brain network structure that may be attributed to development/aging or diseases. The close relation between curvature and network robustness points to the significance of the detected nodes/areas in ensuring robust operation. Thus, this study lays the foundation for a new approach to assess brain network robustness at the nodal level. We argue that the information provided by curvature may be used in combination with other nodal measures for identifying/studying global changes in the brain.

Curvature (averaged across the network) can also provide a global graph measure for brain network robustness quantification. A similar viewpoint has recently been proposed in the context of financial networks and of gene regulatory networks [22, 23, 61]. It is indeed quite interesting that the connection between curvature and the ability of a dynamical process on a network to return to equilibrium after a perturbation (robustness) is observed in such disparate problems (economy, thermodynamics gene regulation and cancer, brain networks). Several other directions may be worthy of investigating in a similar spirit. In particular, studying curvature changes between nodes at the edge level may prove profitable as, in that case, critical changes in interactions between areas in the brain may be easier to detect. We offer these pointers with the caveat that curvature is sensitive to the way connectivity matrices are generated, i.e., curvature is affected by the choice of parcellation scale, tractography algorithms as well as the particular type of diffusion data, e.g., DTI, HARDI, DSI, etc. Therefore, care must be exercised to minimize such possible effects. The present work focused mainly on exploring the concept of node curvature as a measure of robustness of brain structural networks and comparing with alternative measures. In future studies, it will be worthwhile to explore the relevance of curvature to identify areas belonging to the so-called (functional) default mode network, which can be derived from resting-state functional MRI data, as well as to improve the detection of changes related to development/aging or diseases.

## Acknowledgments

This work was partly supported by AFOSR grants FA9550-15-1-0045 and FA9550-17-1-0435, NIH grants P41 EB015894, P30 NS076408, P41 RR013218, P41 EB015902, U24 CA18092401, R01

AG048769, P30 CA008748, NSF Grant ECCS-1509387 and the Fulbright Program.

Data were provided in part by the Human Connectome Project, WU-Minn Consortium (Principal Investigators: David Van Essen and Kamil Ugurbil; 1U54MH091657) funded by the 16 NIH Institutes and Centers that support the NIH Blueprint for Neuroscience Research; and by the McDonnell Center for Systems Neuroscience at Washington University.

Data were also provided in part by the Human Connectome Project, MGH-USC Consortium (Principal Investigators: Bruce R. Rosen, Arthur W. Toga and Van Wedeen; U01MH093765)

funded by the NIH Blueprint Initiative for Neuroscience Research grant; the National Institutes of Health grant P41EB015896; and the Instrumentation Grants S10RR023043, 1S10RR023401, 1S10RR019307.

We thank Drs. Patric Hagmann and Jeffrey Rudie for making the connectivity matrices, respectively published in [26] and [27], freely available via the USC Multimodal Connectivity Database.

## Footnotes

1 The ***topological entropy*** of a graph *G* is the logarithm of the spectral radius of the adjacency matrix *A*, i.e., the logarithm of the maximum of the absolute values of the eigenvalues of *A* [50].

